# Deprivation of dietary fiber in specific-pathogen-free mice promotes susceptibility to the intestinal mucosal pathogen *Citrobacter rodentium*

**DOI:** 10.1101/2021.06.11.448035

**Authors:** Mareike Neumann, Alex Steimle, Erica T. Grant, Mathis Wolter, Amy Parrish, Stéphanie Willieme, Dirk Brenner, Eric C. Martens, Mahesh S. Desai

## Abstract

The change of dietary habits in Western societies, including reduced consumption of fiber, is linked to alterations in gut microbial ecology. Nevertheless, mechanistic connections between diet-induced microbiota changes that affect colonization resistance and enteric pathogen susceptibility are still emerging. We sought to investigate how a diet devoid of soluble plant fibers impacts the structure and function of a conventional gut microbiota in specific-pathogen-free (SPF) mice and how such changes alter susceptibility to a rodent enteric pathogen. We show that absence of dietary fiber intake leads to shifts in the abundances of specific taxa, microbiome-mediated erosion of the colonic mucus barrier, a reduction of intestinal barrier-promoting short-chain fatty acids, and increases in markers of mucosal barrier integrity disruption. Importantly, our results highlight that these low fiber diet-induced changes in the gut microbial ecology collectively contribute to a lethal colitis by the mucosal pathogen *Citrobacter rodentium,* which is used as a mouse model for enteropathogenic and enterohemorrhagic *Escherichia coli* (EPEC and EHEC, respectively). Our study indicates that modern, low-fiber Western diets might make individuals more prone to infection by enteric pathogens via the disruption of mucosal barrier integrity by diet-driven changes in the gut microbiota, illustrating possible implications for EPEC and EHEC infections.

## Introduction

Typical diets in Western societies are characterized by a fiber intake below the recommended amount of 28–38 g per day^1^. Such a reduced consumption of fiber is a possible explanation for the imbalances in gut microbial communities, which have been associated with various diseases such as inflammatory bowel disease (IBD) and colorectal cancer^2^. Furthermore, several studies have shown a link between Western style diets and enteric pathogenic infections in mice^3-5^. Specifically, bacterial metabolites derived from dietary fiber fermentation are known to impact disease severity of enteric pathogenic infections^6^. As shown in mouse models, apart from maintaining immune homeostasis by producing certain metabolites like short-chain fatty acids (SCFAs) from dietary fiber digestion^7,8^, the intestinal microbiome confers colonization resistance against invading enteric pathogens by enhancing mucosal barrier integrity^9^ or by competing for the same nutritional resources^10^. However, knowledge of how deprivation of dietary fiber impacts susceptibility to a mucosal pathogen by the altered gut microbial ecology is still being uncovered.

The murine mucosal pathogen *Citrobacter rodentium* forms attaching and effacing (A/E) lesions on the intestinal mucosa of the host and is often used as a model for human enteropathogenic and enterohemorrhagic *Escherichia coli* (EPEC and EHEC, respectively) infections^11^. Thus, *C. rodentium* allows further study of how deprivation of dietary fiber impacts the complex ecological interactions between the gut microbiome, enteropathogens and the host. This, in turn, can aid in understanding the associations of Western dietary patterns with increased susceptibility to enteric pathogens^12^. The lessons learned from such a model could furthermore eventually inform strategies for designing customized diets to reduce the burden of enteric infections.

We previously showed that, when gnotobiotic mice colonized with a 14-member synthetic human gut microbiome (14SM) are fed a fiber-free diet, a community-wide proliferation of mucus-degrading bacteria causes erosion of the intestinal mucus barrier^13^. This leads to lethal colitis by the rodent mucosal pathogen *C. rodentium*^13^. It was uncertain whether our results from a gnotobiotic mouse model could be observed within specific-pathogen-free (SPF) mice, since the pathogen should face increased colonization resistance from a native, complex gut microbiota. Here, we investigated whether increased susceptibility to *C. rodentium* can be observed in fiber-deprived SPF mice and whether accompanying alterations in the gut homeostasis might contribute to the virulence of this mucosal pathogen.

## Methods

### Ethical statement

The animal experiments were approved by the Animal Welfare Structure of the Luxembourg Institute of Health (protocol number: DII-2017-14) and by the Luxembourgish Ministry of Agriculture, Viticulture and Rural Development (national authorization number: LUPA 2018/09). All experiments were performed according to following the “Règlement grand-ducal du 11 janvier 2013 relatif à la protection des animaux utilisés à des fins scientifiques” based on the “Directive 2010/63/EU” on the protection of animals used for scientific purposes. The mice were housed in a specific-pathogen-free (SPF) facility according to the conditions stated in the recommendations of the Federation of European Laboratory Animal Science Association (FELASA).

### Experimental design and dietary treatment

A total of fifty 6 to 8-week-old, female C57BL/6J mice purchased from Charles Rivers Laboratories (Saint Germain Nuelles, 69210, France) were used for the experiments reported in this study. Mice were randomly housed in the same individually ventilated cage (IVC) rack in groups of up to five animals per cage. Throughout the experiments, all animals were exposed to 12 hours of light daily. Sterile distilled water and diets were provided ad libitum. Upon arrival, all mice were maintained on a standard mouse chow, which we call a fiber-rich (FR) diet, for seven days. After the seven days of the settling time, the cages were randomly allocated to one of the two dietary groups. The first group of five cages was maintained on the FR diet (*n*= 25), while the second group of five cages was switched to a fiber-free (FF) diet (*n* = 25) for 36–40 days. Prior to sacrificing and prior to infection with *C. rodentium*, fecal samples were collected from all fifty mice for various readouts such as lipocalin (Lcn2) quantification (FR, *n*= 25 and FF, *n* = 25), *p*-nitrophenyl glycoside-based enzyme assay (FR, *n* = 4 and FF, *n* = 4), and 16S rRNA gene analyses (FR, *n* = 10 and FF, *n* = 10). After the feeding period, 10 mice (FR, *n* = 5 and FF, *n* = 5) were euthanized by cervical dislocation for mucus layer measurements, histological evaluation, colon length measurements and short-chain fatty acid analyses (cecal contents). Nine animals (FR, *n* = 4 and FF, *n* = 5) were maintained on the two diets for a total of 70 days to control for diet-specific weight alterations. 20 animals were infected with approximately 2 × 10^8^ colony-forming units (CFUs) of kanamycin-resistant, wild-type *Citrobacter rodentium* strain (DBS 120) via intragastric gavage of 0.2 ml of culture (FR, *n* = 10 and FF, *n* = 10; five mice per cage). The same inoculum of *C. rodentium* was used to gavage all mice in both dietary groups. Fecal samples were collected daily following the infection. Mice were scored daily (none, mild, moderate, or severe) for the following criteria: reduced grooming habits resulting in dull coat, weight loss, activity, diarrhea, constipation, anorexia, hunched back position, and dehydration. The same scoring was also used for uninfected animals maintained on the two diets for 70 days. Animals were euthanized if three of the criteria were scored as moderate, one of the physical appearance criterions was scored as severe, or if the animals lost >20% of their initial body weight (measured on the day of infection, before the intragastric gavage of *C. rodentium*).

### Animal diets

The fiber-free (FF) diet was manufactured by SAFE diets (Augy, France), as previously described^13^, and was synthesized according to a modified version of the Harlan TD.08810 diet. The fiber-rich (FR) diet (4.23% crude fiber) is the standard mouse chow in the local animal facility (Special Diets Service; Essex, UK; product code: 801722). The FF diet and the FR diet were both sterilized using 9 kGy gamma irradiation.

### Mucus layer measurements and histological evaluation

After euthanizing the animals by cervical dislocation, the gastrointestinal tracts were dissected, the colons were rapidly separated and fixed in methacarn solution (60% dry methanol, 30% chloroform, 10% glacial acetic acid) and then stored in 100% methanol until further processing. Samples were infiltrated with paraffin in a tissue processor, embedded in paraffin and subjected to either Alcian blue or hematoxylin and eosin (H&E) staining on thin colonic sections (5 µm). Alcian blue-stained sections were used for the mucus layer measurements, as described previously^13^. Samples were deparaffinized in xylene (2x) for 8 min and 5 min, hydrated in 96%, 80%, and 70% ethanol (5 min each) and transferred to distilled water (30 sec). Samples were stained in Alcian blue solution (30 min) and washed under running tap water (2 min), briefly rinsed in distilled water (30 sec), and dehydrated with 96% alcohol (2x) for 1.5 min and 2 min. Samples were transferred to isopropanol (5 min) and treated with xylene (2x) for 5 min each and covered with a coverslip. Partially overlapping tile pictures over the entire colon length were taken by M. N. using a Zeiss Axio Observer Z1 Inverted Phase Contrast Fluorescence Microscope with the Zen software. The mucus layer measurements were performed in a single-blinded fashion by A. P. using ImageJ. Depending on the size of the fecal pellet, between 100– 1300 measurements were performed for each animal and the average mucus layer thickness was calculated for each animal by averaging the measurements. H&E-stained sections were used for blinded histological scoring. Samples were deparaffinized as described above and stained with hematoxylin solution according to Mayer (5 min), washed in distilled water (30 sec) and transferred to running tap water (5 min). Samples were counterstained with 1% aqueous eosin (5 min), washed under running tap water (4 min), dehydrated as described above and covered with a coverslip. Histological scoring was performed with a validated scale^14^ of 0– 3 with “0” representing no inflammation and “3” representing severe inflammation (e.g. infiltration of inflammatory cells, crypt hyperplasia, goblet cell-loss, and distortion of colonic architecture).

### Citrobacter rodentium *growth and fecal quantification*

*C. rodentium* was cultured aerobically from a cryostock on LB-agar plates supplemented with 50 µg/ml kanamycin. A single colony was incubated in Luria-Bertani (LB) broth without antibiotic supplementation at 37 °C for 22 hours. The culture was then diluted in phosphate-buffered saline (PBS) to reach the amount of approximately 2 × 10^8^ CFU per 0.2 ml, which was verified by plating on LB-agar plates supplemented with 50 µg/ml kanamycin. Daily fecal sampling to determine *C. rodentium* CFUs was performed daily and samples were processed immediately. CFU detection continued to be performed daily for FF-fed mice until day 47 post infection, whereas no further CFU detection was performed for FR-fed mice upon pathogen clearance. The samples were weighed, homogenized in 1 ml cold 1xPBS, and 10 µl of the fecal suspensions was plated in serial dilutions on LB-agar plates supplemented with 50 µg/ml kanamycin. The plates were incubated aerobically overnight at 37 °C and the emerging colonies from the appropriate dilutions, ranging from 10^4^ to 10^12^, were counted. For CFU analysis, fresh cecal contents were collected from sacrificed animals and processed immediately, as described above.

### Lipocalin ELISA

A total of *n* = 50 samples (FR, *n* = 25; FF, *n* = 25) were used for this experiment. Lipocalin (Lcn2) concentration was determined for fecal samples stored at –20 °C. Ice-cold PBS with 1% Tween 20 (500 µl) was added to fecal pellets weighing approximately 30 mg, followed by homogenization for 20 min at 4 °C on a thermomixer. Samples were then centrifuged for 10 min at 18 000 × g. Supernatants were stored at –20 °C and Lcn2 detection was performed using the Mouse Lipocalin-2/NGAL DuoSet Elisa, R&D Systems (Bio-Techne), according to the manufacturer’s instructions.

### Colon length measurements

Colons fixed in methacarn solution (see above) were transferred to a histology cassette and stored in 100% methanol. The lengths of colons were measured as described previously^13^.

### Intestinal fatty acid analysis

Flash-frozen cecal contents (30–100 mg) were homogenized using five ceramic beads (1.4 mm) per tube with 500 µl stock solution IS (2-Ethylbutyric acid, 20 mmol/L) per 50 mg of cecal content (VK05 Tough Micro-Organism Lysing Kit). Samples were homogenized at 4 500 **×** g for 30 sec at 10 °C (Precellys24 Homogenizer) and centrifuged for 5 min at 21 000 x g and 4 °C. Further processing of the homogenate and measurements of SCFAs were then performed using high-performance liquid chromatography (HPLC), as previously described^15^.

### p-*Nitrophenyl glycoside-based enzyme assays*

Frozen fecal samples were stored and processed according to an established protocol^16^. The activity of three host mucus targeting bacterial enzymes, β-*N*-acetylglucosaminidase, sulfatase, α-fucosidase, and two dietary plant fiber glycan-targeting bacterial enzymes, α-galactosidase, and β-glucosidase were investigated. Details about bacterial enzymes, their biological substrate and the chemical substrate used for activity detection can be found in **Table 1**.

**Table 1:** Bacterial enzymes and chemical substrates used for activity detection.

| Bacterial enzyme | Biological substrate | Chemical substrate used for activity detection | Supplier |
| --- | --- | --- | --- |
| Sulfatase | Host mucus glycans | Potassium 4-nitrophenyl-sulfate | Sigma Aldrich, Cat#N3877 |
| β-N-Acetyl-glucosaminidase | Host mucus glycans | 4-Nitrophenyl N-acetyl-β-D-glucosaminide | Sigma Aldrich, Cat#N9376 |
| α-Fucosidase | Host mucus glycans | 4-Nitrophenyl α-L-fucopyranoside | Sigma Aldrich, Cat#N3628 |
| α-Galactosidase | Dietary plant fiber glycans | 4-Nitrophenyl α-D-galactopyranoside | Sigma Aldrich, Cat#N0877 |
| β-Glucosidase | Dietary plant fiber glycans | 4-Nitrophenyl β-D-glucopyranoside | Sigma Aldrich, Cat#N7006 |

### Illumina sequencing of 16S rRNA genes and data analysis

Fecal samples were stored at –20 C° until bacterial DNA extraction, as previously described and library preparation were also performed as described previously^13^. The cited protocol is a modified version of the protocol by Kozich et al. in which the V4 region of the 16S rRNA gene is amplified using dual-index primers^17^. Raw sequences have been deposited in the European Nucleotide Archive (ENA) under the study accession number PRJEB44016. The raw sequences were processed using the program mothur (v1.33.3)^18^ according to the MiSeq SOP published on the mothur website^17,19^. Probabilistic modelling to cluster the microbial communities in metacommunities was performed in mothur based on the Dirichlet multinomial mixtures method^20^. Alignment and classification of the sequences was done using the SILVA reference database (release 132)^21^. Chimeric sequences were removed using the VSEARCH tool. Microbial abundance figures were generated using the following packages in RStudio 4.0.2: phyloseq 1.34.0, ggplot2 3.3.3, vegan 2.5.7, forcats 0.5.1. Alpha diversity analyses were performed on unfiltered data. The PCoA plot is based on Bray-Curtis dissimilarity index, after filtering out taxa not seen at least 10 times in at least 20% of the samples. The spatial means of the fiber-rich (FR) diet-fed mice and fiber-free (FF) diet-fed mice clusters were significantly different by PERMANOVA using adonis (*p* = 0.001), although a significant betadisper test (*p*= 0.009) means that this finding could be an artifact of the heterogenous dispersions. The heatmap was generated based on the top 50 significant OTUs determined using DESeq2; all OTUs shown are significant based on an adjusted *p*-value<0.01. The heatmap shows log2 fold-change for data rarefied to the minimum library size and filtered (minimum 10 counts in 20% of samples) displaying the fold-changes in abundance of each OTU, relative to the average abundance of that OTU in the comparator dietary group. Samples where the read count is zero are presented as infinitely negative values. In cases where all samples in a group had zero read counts, all samples with a non-zero value in the comparator group are then infinitely positive relative to the other group's zero average count. Adjusted *p*-values were obtained using the Benjamini-Hochberg procedure to correct for multiple comparisons.

## Results and discussion

Six to eight-week-old, age-matched specific-pathogen-free (SPF) female C57BL/6J mice were maintained on a mouse chow, which, with a fiber content of 4.23% crude fiber, we consider a fiber-rich (FR) diet. Ten animals were switched to a fiber-free (FF) diet and were continuously fed the FF diet for a 36–40 days “feeding period” (**Fig 1a**). As controls, the remaining C57BL/6J mice were maintained on the FR diet. The FR diet contained naturally milled food ingredients with crude plant fiber sources, such as corn, soybean, and wheat, whereas the isocaloric FF diet lacked such plant polysaccharides and instead contained increased levels of glucose^13^. Throughout the feeding period, the FF-fed mice did not exhibit any physiological irregularities (data not shown) or weight loss.

**Figure 1:**
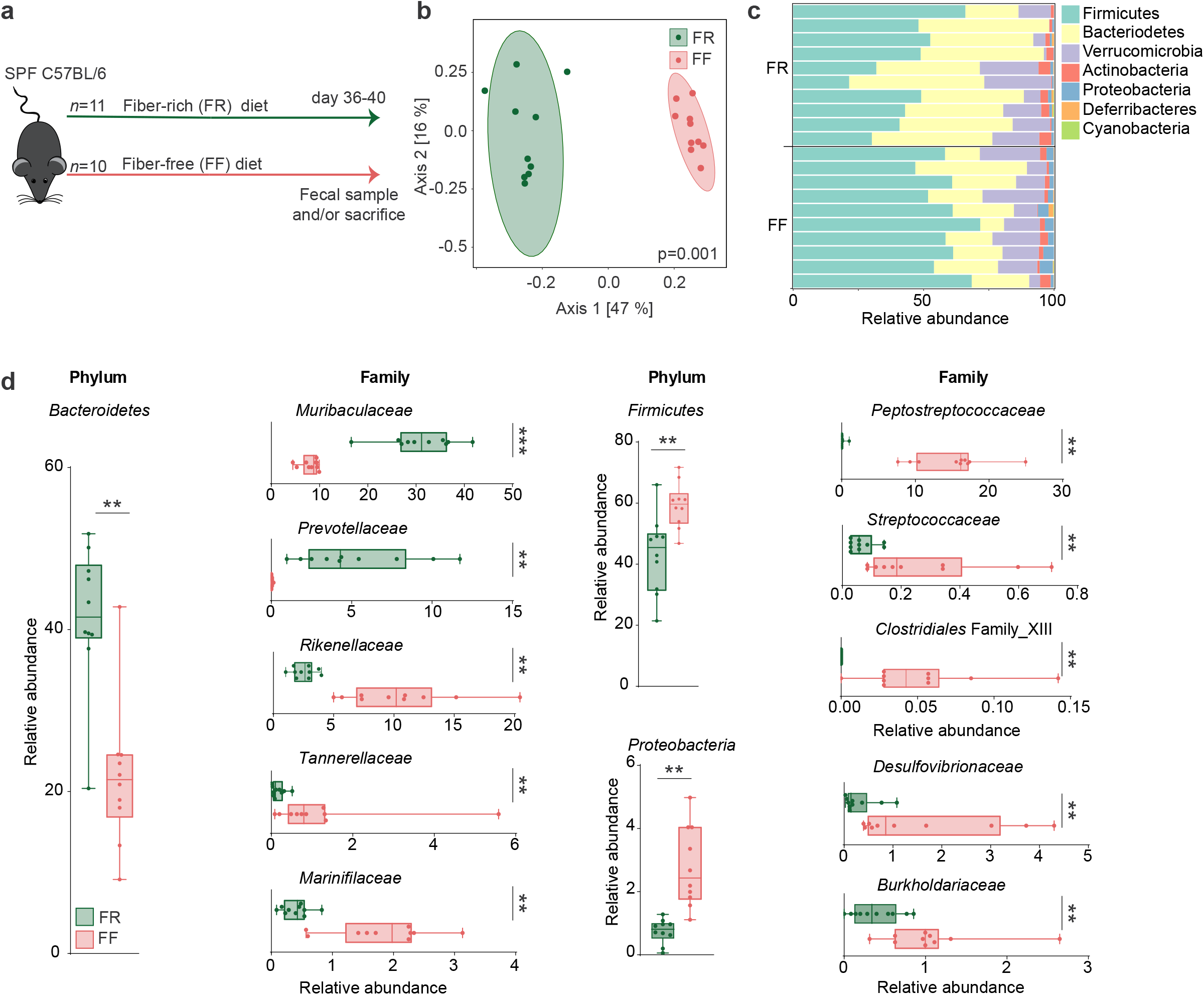
Fiber deprivation in mice harboring a complex microbiome results in changes in bacterial families. Green: FR-fed mice; Red: FF-fed mice. * = *p*<0.05; ** = *p*<0.01; *** = *p*<0.001. **(a)** Experimental schedule. 6 to 8 week-old female specific-pathogen-free (SPF) C57BL/6J mice were continued on a fiber-rich (FR) diet or switched to a fiber-free (FF) diet for a feeding period of 36–40 days. At the end of the 36–40 days feeding period, mice were sacrificed for various readouts. FR, *n* = 11 and FF, *n* = 10. **(b)** Principal coordinates analysis (PCoA) of the fecal microbial communities of FR and FF mice after 36 days of feeding, based on Bray-Curtis dissimilarity index. Ellipses represent 95% confidence interval for each group. The difference in spatial means of FR and FF diet clusters was significant by PERMANOVA testing using adonis (*p* = 0.001), however, the dispersions were also significantly different by betadisper (*p* = 0.009). FR, *n* = 10 and FF, *n* = 10. **(c)** Barplot of relative microbial abundances of the bacterial phyla after 36 days of feeding among FR-(*n* = 10) and FF-fed (*n* = 10) mice. **(d)** Boxplots displaying significant changes in the relative abundance (%) of *Bacteroidetes*, *Firmicutes* and *Proteobacteria* between FR-(*n* = 10) and FF-fed (*n* = 10) mice. Boxplots display significant changes in relative abundance (%) of bacterial families between FR- and FF-fed animals, sorted by their corresponding phylum. Whiskers display minimum and maximum. Mann-Whitney test, two-tailed. Adjusted p-values were obtained using the Benjamini-Hochberg procedure to correct for multiple comparisons.

### Fiber deprivation leads to an increase in potential mucus-degrading bacterial taxa in SPF mice

After the feeding period, 16S rRNA gene sequencing (Illumina platform) was performed using DNA extracted from the mouse fecal samples. The results were compared to the FR-fed mice to evaluate changes in microbiome composition in response to diets with different fiber content (**Fig. 1b–d)**. Alpha diversity between groups was not significantly different using the Chao 1 richness estimate or by Shannon and inverse Simpson indices, which account for the evenness of the detected taxa (data not shown). Principal Coordinates Analysis (PCoA) based on Bray-Curtis dissimilarity (**Fig. 1b**) revealed significant diet-specific clustering by PERMANOVA (*p* = 0.001). Although the dispersion was significantly higher among the FR-fed mice (*p* = 0.009), it is reasonable to conclude that the spatial medians are indeed significantly different since the design is balanced^22^. On the phylum-level, *Proteobacteria* and *Firmicutes* were significantly higher among FF-fed mice, whereas *Bacteroidetes* was enriched in FR-fed mice (**Fig. 1c**). Family-level analysis revealed heterogeneous changes within the *Bacteroidetes* phylum, as *Muribaculaceae* and *Prevotellaceae* were enriched in FR-fed mice, but *Rikenellaceae*, *Marinifilaceae*, and *Tannerellaceae* were higher among FF-fed mice (**Fig. 1d**). *Firmicutes* belonging to *Peptostreptococcaceae*, *Streptococcaceae*, and *Clostridiales* Family XIII, were higher among FF-fed mice, as were *Proteobacteria* belonging to *Burkholderiaceae* and *Desulfovibrionaceae* families (**Fig. 1d**). An expansion of *Proteobacteria* has been associated with intestinal epithelial dysfunction^23^. *Peptostreptococcaceae* have been found to be enriched in patients with colorectal cancer^24^ and has been associated with a decrease in mucus layer thickness^25^. Furthermore, increased levels of *Bacteroidetes* over *Firmicutes*, as shown in FR-fed animals (**Fig. 1d**), has been previously linked with a decreased *C. rodentium* susceptibility^26^. Analysis of log2 fold-change on an OTU level using DESeq2 reveals discrete changes between the dietary conditions (**Fig. 2**). We exclude a cage effect based on the Dirichlet multinomial mixtures method^20^, which identified only two microbial communities clusters that correspond to the diet. These clusters are evident in the phylogenetic tree of the heatmap, depicting samples from individual animals and their corresponding cages (**Fig. 2**). An upregulation of specific OTUs belonging to the bacterial genera *Lachnospiraeceae, Ruminococcaceae,* and *Oscillibacter* could be detected in fiber-deprived animals compared to the control group (**Fig. 2**). An increased abundance of *Lachnospiraeceae* spp. has previously been reported to be associated with the increased susceptibility to adherent-invasive *Escherichia coli* (AIEC) infection^27^. Furthermore, *Ruminococcaceae* spp.^28^, *Lachnospiraceae* spp., *Clostridiales* spp., *Desulfovibrionales* spp., and *Oscillibacter* spp. have been shown to be correlated with the production of branched-chain fatty acids (BCFAs), such as valeric, isovaleric and isobutyric acid^29^.

**Figure 2:**
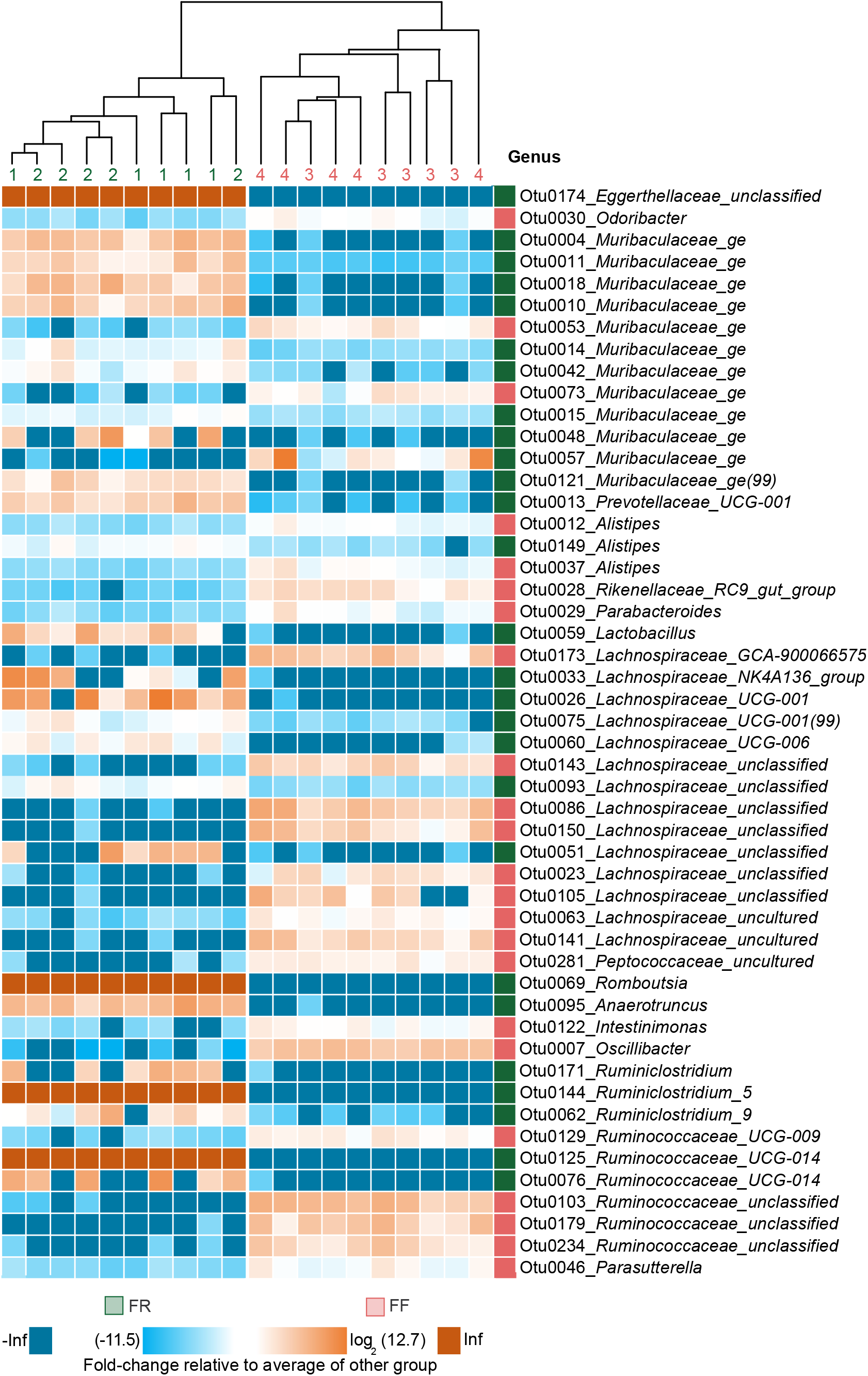
Fiber deprivation in mice harboring a complex microbiome results in changes in bacterial families with no detectable cage effect. Phylogenetic tree displaying clustering by cage (1–4). Green numbers indicate FR-fed mice and red numbers indicate FF-fed mice. FR, *n* = 10, FF, *n* = 10 with *n* = 5 animals per cage. Heatmap displaying fold-changes in abundance of top 50 operational taxonomic units (OTUs) that were significantly different between FR- and FF-fed mice using DESeq2, based on the adjusted *p*-value; FR, *n* = 10, FF, *n* = 10. The log_2_-transformed fold-change of each OTU (row) in each sample (columns) is calculated relative to the average abundance of the other group. OTUs labelled with green squares are enriched in FR-fed mice, while OTUs labelled with red squares are enriched in FF-fed mice. Adjusted *p*-values were obtained using the Benjamini-Hochberg procedure to correct for multiple comparisons.

In line with these and previous results, we investigated whether dietary fiber deprivation resulted in altered intestinal fatty-acid concentrations by measuring levels of SCFAs (acetate, propionate, and butyrate) and BCFAs (valerate, isovalerate, and isobutyrate) in cecal contents of FR-fed and FF-fed mice at the end of the feeding period. We detected significantly increased levels of all SCFAs in FR-fed mice compared to FF-fed mice, while amounts of all detected BCFAs were significantly lower compared to fiber-deprived mice (**Fig. 3a**). While SCFAs are generated largely by the microbial fermentation of dietary fibers^30^, intestinal BCFAs are mainly the result of bacteria-mediated dietary protein metabolism^31^.

**Figure 3:**
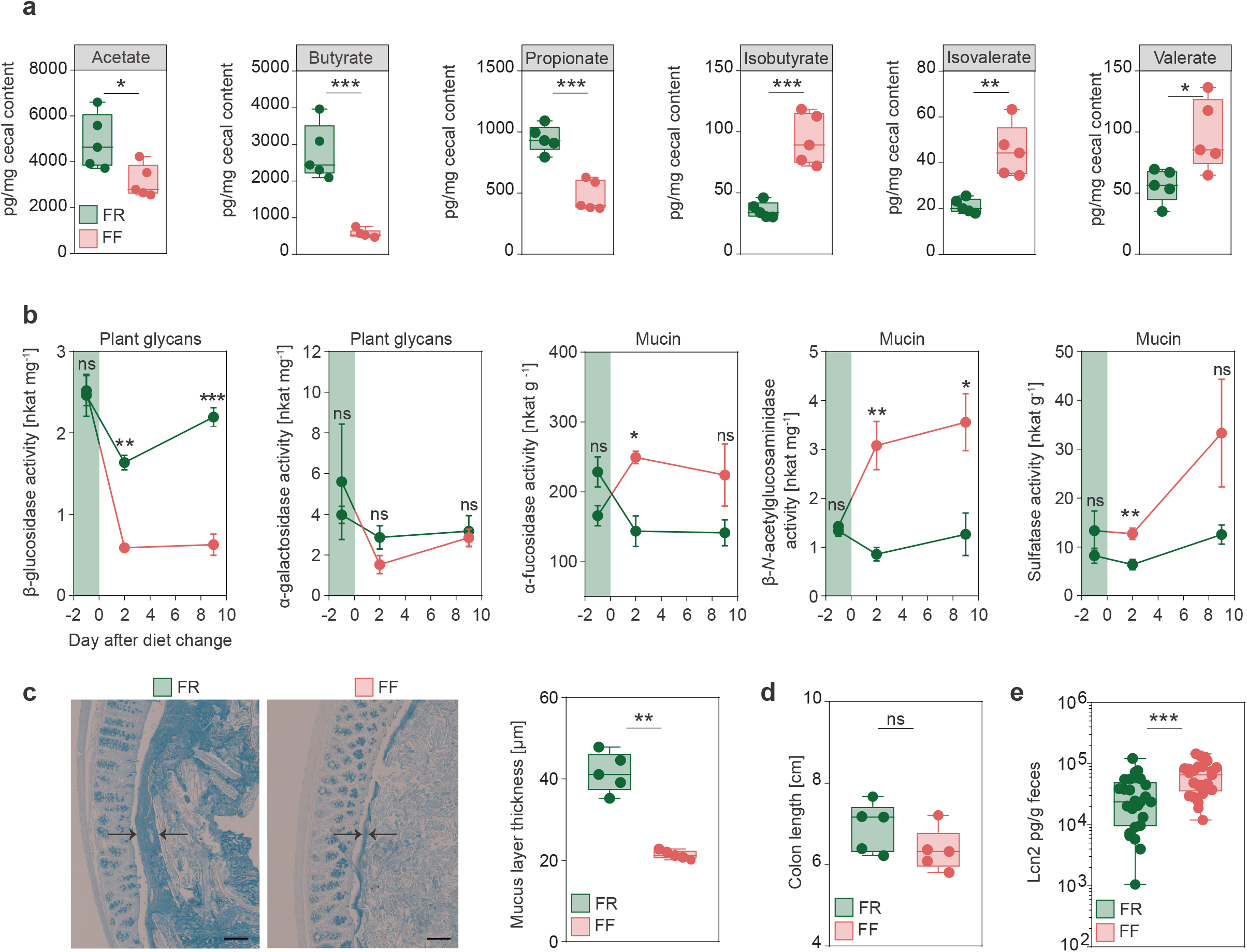
Fiber-deprivation in mice harboring a complex microbiome results in changes in bacterial enzyme activity and levels of mucosal barrier integrity markers. Green: FR-fed mice; Red: FF-fed mice. * = *p*<0.05; ** = *p*<0.01; *** = *p*<0.001. **(a)** SCFA concentrations in cecal contents in pg SCFA per mg cecal content after 40 days of feeding in FR-(*n* = 5) and FF-fed (*n* = 5) animals. Unpaired t-test, two-tailed. **(b)** Results for *p*-nitrophenyl glycoside-based enzyme assay from fecal samples of FR-fed (*n* = 4) and FF-fed (*n* = 4) animals. Evaluation of five different bacterial enzymes. Plant glycans: β-glucosidase, α-galactosidase. Mucin: α-fucosidase, β-*N*-acetylglucosaminidase, and sulfatase at three different time points. Day 1 before diet switch to FF diet, day 2 after diet switch to FF diet and day 9 after diet switch to FF diet. Green area indicates FR feeding period before diet switch to FF diet. Error bars represent SEM. Unpaired t-test, two-tailed. **(c)** Left panel: Alcian blue-stained 5 µm thin sections of the colonic mucus layer of FR-fed and FF-fed animals. Black arrows indicate thickness of the colonic mucus layer. Scale bar = 100 µm. Right panel: Statistical analysis of mucus layer measurements of FR-fed (*n* = 5) and FF-fed (*n* = 5) animals. Mann-Whitney test, two-tailed. **(d)** Colon length in FR-(*n* = 5) and FF-fed (*n* = 5) mice after 40 days of feeding. Mann-Whitney test, two-tailed. **(e)** Levels of lipocalin in fecal supernatant, normalized on fecal weight for FR-(*n* = 25) and FF-fed (*n* = 25) animals after 36–40 days of feeding. Mann-Whitney test, two tailed.

Among other microbial protein fermentation products, such as phenol, biogenic amines, hydrogen sulfide and p-cresol^32^, BCFAs are associated with an increased colonic epithelial cell permeability *in vitro*^33^, whereas SCFAs are reported to maintain mucosal barrier integrity by increasing mucin-2 glycoprotein (MUC2) expression^34^, enhancing colonic tight-junction assembly^35^, and stimulating antimicrobial peptide secretion^36^. Furthermore, changes in intestinal butyrate concentration is known to affect susceptibility of *C. rodentium*^37^ and oral administration of butyrate ameliorates *C. rodentium* induced inflammation via enhanced IL-22 production^38^.

Additionally, specific species within the *Rikenellaceae* (*Alistipes*)^39,40^, *Lachnospiraceae*^41,42^ and *Muribaculaceae*^40^ families are thought to possess the capacity to degrade mucus. Thus, in line with our 14SM model, our results suggest that the deprivation of dietary fiber may also increase mucus-degrading populations in the SPF mice containing their native gut microbiota. An increase in *Desulfovibrionaceae* family (**Fig. 1d**) is consistent with the significant increase of *Desulfovibrio piger* (family: *Desulfovibrionaceae*), which was previously observed in fiber-deprived 14SM-colonized gnotobiotic mice^13^. An increase in these sulfate-reducing bacteria indicates availability of increased sulfate groups that are used as an electron acceptor by this group of bacteria. A possible source of this increased sulfate could be the terminal sulfate groups on the colonic mucins^10^.

### A dietary fiber-deprived gut microbiota erodes the colonic mucus layer in SPF mice

Given the relevance of mucus erosion on pathogen-susceptibility of fiber-deprived mice in a gnotobiotic mouse model^13^, we evaluated whether the increase in these potentially mucus-degrading taxa resulted in the increased activities of microbial enzymes that are involved in the degradation of host-secreted colonic mucus. We studied activities of five different enzymes in the mouse fecal samples before, and after a switch to the FF diet. Our results show a significant increase in the activities of three microbial enzymes, namely α-fucosidase, β-*N*-acetylglucosaminidase and sulfatase (**Fig 3b**), that are involved in colonic mucus degradation. Conversely, the activity of β-glucosidase, an enzyme involved in the degradation of plant glycans, was significantly decreased in the FF-fed animals.

To assess the impact of the increased activities of these mucus-specific enzymes on the mucus layer, we performed measurements of the colonic mucus layer thickness, using fixed colonic tissue samples stained with Alcian blue^13^. The mucus measurements revealed significantly decreased mucus layer thickness in fiber-deprived mice compared to the FR-fed mice (**Fig. 3c**). Taken together, decreased SCFA levels and reduced mucus layer thickness suggest impairment of the intestinal mucosal barrier in fiber-deprived mice.

In order to evaluate inflammatory reactions in the colonic tissue in response to the potential mucosal barrier impairment in the FF-fed mice, we measured colon lengths and the levels of lipocalin (Lcn2), a sensitive marker capable of detecting low-grade inflammation, in fecal contents of the FR- and FF-fed mice after the feeding period. There was a trend towards reduced colon lengths in the FF-fed animals compared to the FR-fed animals (**Fig. 3d**), athough this was not statistically significant. Fecal Lcn2 levels, however, were significantly increased in fiber-deprived mice (**Fig. 3e**), indicating an increased inflammatory tone in FF-fed mice without pronounced colonic inflammation. Furthermore, histopathological evaluation^14^ of colonic hematoxylin/eosin-stained sections did not reveal significant differences between FR- and FF-fed mice (histology data not shown). These observations were in line with the previous observation in our previous gnotobiotic model^13^ and prompted us to investigate whether the diet-induced alterations in the microbiome composition and its associated effects on the markers of mucosal barrier integrity impacted the susceptibility of fiber-deprived mice to infection with *C. rodentium*.

### *Deprivation of dietary fiber in SPF mice promotes susceptibility to* C. rodentium

To test our hypothesis, we infected FF-fed and control FR-fed mice with *C. rodentium.* Weight and general appearance were monitored daily for up to 47 days post infection (dpi), and fecal *C. rodentium* (*Cr*) colony forming units (CFUs) were assessed until all animals cleared the pathogen or had to be euthanized (**Fig. 4a**). FR and FF animals without infection showed no significant weight differences in two dietary groups for up to 70 days after switch to the FF diet (**Fig. 4b**). The FF-fed animals infected with *Cr* demonstrated significantly higher weight loss compared to the FR-fed animals, with several animals requiring euthanasia owing to >20% weight loss (**Fig. 4c**). In line with the observed weight loss, survival curves of infected animals fed the two different diets showed only a 40% survival rate in FF-fed mice, whereas 100% of the mice in the FR-fed group survived (**Fig. 4d**).

**Figure 4:**
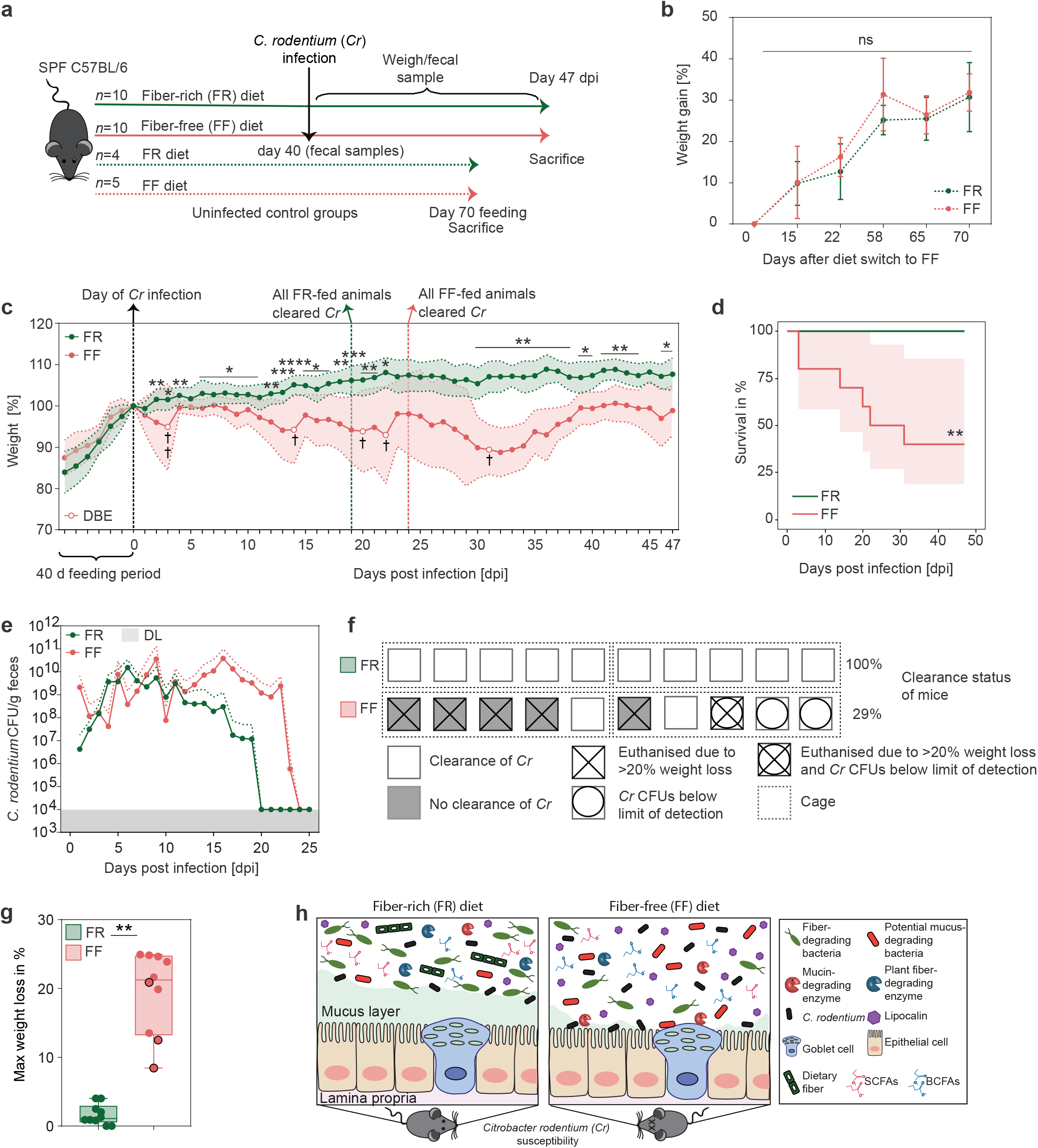
Fiber-deprivation in mice is associated with a higher susceptibility to *C. rodentium* infection. Green: FR-fed mice; Red: FF-fed mice. * = *p*<0.05; ** = *p*<0.01; *** = *p*<0.001; **** = *p*<0.0001. **a)** Experimental schedule. Animals were fed the FR diet or the FF diet (FR, *n* = 10 and FF, *n* = 10) for 40 days and infected via intragastric gavage with 2 × 10^8^ colony forming units (CFU) in 0.2 ml culture of *C. rodentium* (*Cr*). Fecal samples were collected before the infection and used for various readouts. Animals were monitored daily for 47 days post infection (dpi). Mice were euthanized when they lost 20% of the body weight compared to the initial day of infection. Dotted red and green arrow: animals (FR, *n* = 4 and FF, *n* = 5) were fed the respective diets as uninfected control groups for 70 days. **(b)** Mean weight in percent of FR-fed and FF-fed animals over a time period of 70 days after the diet switch. FR: *n* = 4 and FF: *n* = 5. Green and red error bars represent SD. ns = not significant. T-test, two-tailed. **(c)** Left of zero: Mean weight as a percent of reference weight to the day of diet switch of FR-fed (*n* = 15; 5 animals used for earlier readouts without infection) and FF-fed (*n*= 15; 5 animals used for earlier readouts without infection) animals during the 40-days feeding period. Dotted black line at d0: day of *Cr* infection and reference weight. Right of zero: Mean weight as a percent of reference weight (d0) in FR-fed (*n* = 10) and FF-fed (*n* = 10) animals after the infection with *Cr*. Dotted green or red line: all FR- or FF-fed animals cleared *Cr*, respectively. White circles indicate time points where animals had to be euthanized due to weight loss beyond 20%, compared to the reference weight (d0). Black cross indicates number of animals that had to be euthanized. DBE = death before experimental endpoint. Red and green areas represent SD. Mann-Whitney test, two tailed. **(d)** Cumulative number of individuals at risk over time among FR-(*n* = 10) and FF-fed (*n* = 10) animals infected with *Cr*. Log-rank test. Red area represents confidence interval. **(e)** *Cr* CFUs per gram fecal content of FR-(*n* = 10) and FF-fed (*n* = 10) animals. For days where CFUs were below the detection limit of 10^4^ CFUs, the detection limit of 10^4^ CFUs was used. No CFUs values were included if no fecal samples could be obtained or if an animals had to be euthanized. Red and green dotted lines represent SD. DL= detection limit of 10^4^ CFUs. **(f)** Graphic representation of the clearance rates of *Cr* susceptible animals, FR, *n* = 10 and FF, *n* = 10. Animals and treatment groups sorted by their caging (5 animals/cage) (dotted lines) and dietary experimental groups. Each symbol represents one mouse with the respective clearance status and *Cr* susceptibility. **(g)** Maximal weight loss of each individual animal in percent (FR, *n* = 10 and FF, *n* = 10). Maximal weight loss is displayed in percent of the day during the experiment where the highest weight loss was detected. Black circles: Animals displaying *Cr* CFUs below limit of detection throughout the experiment. Wilcoxon matched-pairs signed rank test. Two-tailed. **(h)** Graphical summary on diet-driven changes within gut homeostasis on *Cr* susceptibility. Left: Homeostatic conditions in FR-fed mice. Right: Changes in dietary fiber-deprived mice within gut homeostasis. Far right: Legend.

Moreover, FF-fed mice showed overall delayed *Cr* clearing capacities and elevated CFUs in fecal samples until day 23 dpi (**Fig. 4e**). While the FR-fed mice showed rapidly decreasing CFUs after 11 dpi and all mice of this group cleared the pathogen completely by 19 dpi (**Fig. 4e**), the FF-fed mice took longer (24 dpi) to clear the pathogen. Even though three FF animals failed to show detectable CFUs, one out of these three FF animals had to be euthanized on 31 dpi due to weight loss (**Fig. 4c)**, indicating that fiber deprivation acts in favor of a lethal *Cr* phenotype even when CFUs are below the detection limit. These results suggest that dietary fiber deprivation is associated with a delayed *Cr* clearance (**Fig. 4e**) as well as reduced rates of clearance (**Fig. 4f**) and an overall higher weight loss (**Fig. 4g**). The infection by *Cr* is usually self-limiting in healthy mice because the commensal gut bacteria are able to outcompete the pathogen by competing for nutritional resources, such as monosaccharides^43,44^.

A lethal infection phenotype like the one we observed in the FF-fed wild type mice has typically been observed in mice lacking essential genes that protect from *Cr* clearance and invasion, such as *Muc2* ^45^ or genes encoding the cytokines IL-10 and IL-22^46,47^. Our previous^13^ and current studies show that such a lethal phenotype could be generated in wild-type mice merely by eliminating crude plant fibers from the diet. Given that *Muc2*^-/-^ mice exhibit a similar lethal phenotype to *Cr*^45^, it is plausible that the fiber-deprived gut microbiota-led erosion of the mucus layer contributes to the pathogenicity. Nevertheless, germ-free mice, which naturally possess a thinner mucus layer, on either the FR and FF diets, do not exhibit a lethal colitis following the *Cr* infection^13^. Thus, together with our previous study^13^, the current study further supports the role of a fiber-deprived, mucus-eroding gut microbiota in driving susceptibility to *Cr*. Notably, no systemic spread of *Cr* into other organs, such as liver or spleen, could be detected in FF-fed mice that had to be euthanized due to >20% weight loss. These results indicate a local impact of *Cr* in the cecum and colon, possibly driven by differences in the microbiota composition and microbial mucus erosion, that drives the lethal phenotype.

## Conclusion

Our results indicate a crucial role of fiber intake on the maintenance of mucosal barrier integrity governed by the microbial ecology of the native, complex mouse microbiome, which plays an important role in protecting against infection with a mucosal pathogen. Different microbiota compositions, in mice from distinct mouse facilities, are responsible for a variable colitis phenotype to *Cr* infection^37^. Here, we show that mice possessing the same starting microbiota, when fed a fiber-deprived diet, undergo a change in the composition and function of the gut microbiota, which collectively contributes to a lethal phenotype to *Cr* (**Fig. 4h**).

A recent study showed that the Western-style diet (WSD) as compared to a standard grain-based chow (GBC) impedes the initial colonization of *Cr*, although WSD leads to alterations in the gut microbiota in such a way that *Cr* cannot be outcompeted by the microbiota^48^; hence, *Cr* in the mice fed WSD cannot be cleared or the clearance is delayed. Here, we observed a similarly delayed clearance of *Cr* in mice fed the FF diet. However, the difference in the phenotype we observe in the FF diet and the one observed in the WSD-fed mice could be rooted in important differences in the composition of these two diets. While the WSD contains nearly twice the amount of %kcal derived from fat as compared to the FF diet (60% vs. 34.1%), the FF diet contains more than twice the calories derived from carbohydrates as compared to the WSD (42.4% vs 20%). Other reasons for differences in the *Cr* infection phenotypes observed with the FF diet and the WSD could be related to the differences in the native mouse gut microbiota and/or differences in the initial pathogen CFU gavaged in the mice.

Compositional differences between the FR and FF diets, other than the fiber content such as increased amounts of simple sugar in the FF diet, raise a caveat in directly associating the lethal phenotype observed in FF-fed mice with the absence of plant fiber. Although we cannot rule out such potential effects of other compositional differences between the FR and FF diets, using our 14M community, we have documented that the two diets lead to a major impact on the fiber-versus mucin-utilization dynamics of the gut microbiota^13^. Furthermore, our current data clearly show that the FF diet leads to enhanced mucin degradation by the gut microbiota, which is a result of fiber deprivation. Thus, it is likely that the absence of plant fiber in the FF diet plays a crucial role on the lethal phenotype observed in this study.

Further investigations are needed to determine specific dietary strategies and/or usage of various reduced microbial communities in gnotobiotic mouse models to understand which specific bacterial taxa, their associated bacterial enzymes or microbial cross-talk with the host immune system modulate susceptibility to *Cr* infection. A follow-up study to the current one needs to utilize various reduced synthetic communities, such as removing some or all four mucus degraders from our 14SM model^13^, to better understand the mechanisms of how mucus-degrading bacteria possibly contribute to the lethal phenotype of *Cr* under fiber deprivation. Removing specific pathogen-promoting and mucus-degrading bacterial species could aid to understand if targeted microbiome modulations could be a tool to prevent severe disease outcomes. Such future studies would also fascilitate the understanding whether the changes observed here (mucus reduction, changes in microbial metabolites, and elevated inflammation) independently contribute to the lethal phenotype of *Cr* or whether a combination of such changes is essential in order to drive susceptibility to *Cr*.

Although the burden of intestinal pathogenic infections is increased and is often more lethal in developing countries as compared to westernized countries^49^, further studies elaborating the connection between a Western style diet and enteric infections could help to prevent associated diseases such as irritable bowel syndrome^50^ and tackle emerging antibiotic overuse^51^. Our findings underline the importance to increased dietary fiber consumption in Western style diets in order to strengthen colonization resistance and barrier integrity within the host against possible enteric infections by EPEC and EHEC.

## Declarations

### Availability of data and materials

Most of the data generated or analysed during this study are included in this article. Please contact author for data requests.

### Competing interests

The authors declare that they have no competing interests.

## Acknowledgements

Work in the authors’ laboratory was supported by the following grants to M.S.D.: Luxembourg National Research Fund (FNR) CORE grants (C15/BM/10318186 and C18/BM/12585940). M.N. and A.P. and were supported by the FNR AFR bilateral grant (15/11228353) and FNR AFR individual grant (11602973), respectively. E.T.G. was supported by the FNR PRIDE grant (17/11823097) and the Fondation du Pélican de Mie et Pierre Hippert-Faber under the aegis of the Fondation de Luxembourg. D.B. is supported by the FNR-ATTRACT program (A14/BM/7632103). E.C.M. was supported by grants from the US National Institutes of Health (DK118024 and DK125445).

## Authors’ contributions

Conceptualization M.N., D.B., E.C.M., and M.S.D.; Experiments, M.N., M.W., S.W., and A.P.; Investigation, M.N., A.S., E.T.G., and M.S.D.; Resources, M.N., S.W., E.T.G, M.W., A.P., A.S., E.C.M., and M.S.D.; Writing – Original Draft, M.N., A.S., and M.S.D; Writing – Review & Editing, M.N., A.S., E.T.G., A.P., M.W., E.C.M. and M.S.D.; Supervision, M.S.D.; Funding Acquisition, M.S.D.

## Notes

### Competing Interest Statement

The authors have declared no competing interest.

